# Novel phosphatidylinositol flippases contribute to phosphoinositide homeostasis in the plasma membrane

**DOI:** 10.1101/2023.07.03.547512

**Authors:** Yumeka Muranaka, Ryo Shigetomi, Yugo Iwasaki, Asuka Hamamoto, Kazuhisa Nakayama, Hiroyuki Takatsu, Hye-Won Shin

## Abstract

Phosphatidylinositol is a precursor of various phosphoinositides, which play crucial roles in intracellular signaling and membrane dynamics and have impact on diverse aspects of cell physiology. Phosphoinositide synthesis and turnover occur in the cytoplasmic leaflet of the organellar and plasma membranes. P4-ATPases (lipid flippases) are responsible for translocating membrane lipids from the exoplasmic (luminal) to the cytoplasmic leaflet, thereby regulating membrane asymmetry. However, the mechanism underlying phosphatidylinositol translocation across cellular membranes remains elusive. Here, we discovered that the phosphatidylcholine flippases ATP8B1, ATP8B2, and ATP10A can also translocate phosphatidylinositol at the plasma membrane. To explore the function of these phosphatidylinositol flippases, we used cells depleted of CDC50A, a protein necessary for P4-ATPase function. Upon activation of the Gq-coupled receptor, depletion of phosphatidylinositol 4,5-bisphosphate [PtdIns(4,5)P_2_] was accelerated in *CDC50A* knockout cells compared with control cells, suggesting a decrease in PtdIns4,5P_2_ levels within the plasma membrane of the knockout cells. These findings highlight the pivotal role of P4-ATPases in maintaining phosphoinositide homeostasis and suggest a mechanism for enrichment of phosphatidylinositol in the cytoplasmic leaflet of the plasma membrane.

## Introduction

Biological membranes exhibit transbilayer lipid asymmetry. In mammalian cells, phosphatidylserine (PtdSer), phosphatidylethanolamine (PtdEtn), and phosphatidylinositol (PtdIns) are abundant in the cytoplasmic leaflet, whereas phosphatidylcholine (PtdCho) and sphingomyelin (SM) are enriched in the exoplasmic leaflet of the plasma membrane (Devaux, 1991; Murate *et al*, 2015; Zachowski, 1993). Phospholipids are mostly synthesized on the cytoplasmic side of the endoplasmic reticulum (ER). In the post-Golgi compartments, phospholipids are distributed asymmetrically through the action of ATP-dependent lipid flippases and floppases, which translocate lipids from the luminal/exoplasmic to the cytoplasmic leaflet and vice versa, respectively. P4-ATPases translocate membrane lipids from the exoplasmic/luminal to the cytoplasmic leaflets of cellular membranes (Andersen *et al*, 2016; Lopez-Marques *et al*, 2020; Shin & Takatsu, 2019; Vance, 2015).

In previous studies, we showed that the human P4-ATPases ATP11A and ATP11C flip NBD-labeled PtdSer (NBD-PtdSer) and NBD-PtdEtn; ATP8B1, ATP8B2, and ATP10A flip NBD-PtdCho; and ATP10D flips NBD-glucosylceramide at the plasma membrane (Naito *et al*, 2015; Roland *et al*, 2019; Takada *et al*, 2015; Takatsu *et al*, 2014). These P4-ATPases interact with CDC50A, which is required for their transport from the ER to the plasma membrane (Bryde *et al*, 2010; Naito *et al*., 2015; Takatsu *et al*, 2011; van der Velden *et al*, 2010b). Mutations in the human *ATP8B1* gene cause progressive familial intrahepatic cholestasis 1 (PFIC1) (Folmer *et al*, 2009; Paulusma *et al*, 2006). Some PFIC ATP8B1 mutant variants fail to flip PtdCho, indicating that PtdCho flipping activity at the bile canaliculi is critical for proper bile excretion into the liver (Takatsu *et al*., 2014). Increased PtdCho flipping activity at the plasma membrane due to exogenous expression of ATP10A in HeLa cells induces the formation of inward membrane curvature and facilitates endocytosis (Takada *et al*, 2018). Moreover, exogenous expression of ATP10A results in cell shape changes and delays cell adhesion and spreading on the extracellular matrix (Miyano *et al*, 2016; Naito *et al*., 2015), indicating the importance of the transbilayer balance of phospholipid mass and/or species. The mouse *Atp10a* gene is linked with diet-induced obesity and type-2 diabetes phenotypes (Dhar *et al*, 2004), and genome wide association studies of nondiabetic patients identified two single-nucleotide polymorphisms in the *ATP10A* gene that are associated with insulin resistance (Irvin *et al*, 2011). ATP8B2 is ubiquitously expressed and seems to be abundant in the brain (Wang *et al*, 2018a), but its physiological function is unknown.

Phosphoinositides are signaling lipids in the cytoplasmic leaflet. They are produced from PtdIns and other phosphoinositides by various kinases and phosphatases (Balla, 2013; Posor *et al*, 2022; Schink *et al*, 2016). Enzymatic interconversion creates a complex network and the cellular levels of phosphoinositides are interdependent. While PtdIns constitutes approximately 10-20% of total phospholipids, the most abundant phosphoinositides, PtdIns 4,5-bisphosphate [PtdIns(4,5)P_2_] and PtdIns 4-phosphate [PtdIns(4)P], account for only 0.2-1%. The other phosphoinositides are present in even smaller amounts (Balla, 2013). The spatiotemporal control of phosphoinositide kinase and phosphatase activity accounts for distinct phosphoinositides becoming enriched in the plasma membrane or subcellular organelles. For instance, PtdIns(4,5)P_2_ is mostly abundant in the plasma membrane, whereas PtdIns(4)P is enriched in the *trans*-Golgi network and PtdIns(3)P in early endosomes (Balla, 2013; Posor *et al*., 2022; Schink *et al*., 2016). Phosphoinositides were originally discovered as rare phospholipid species that undergo turnover upon treatment with cholinergic drug (Hokin & Hokin, 1953). Upon activation of phospholipase C (PLC) by Gq-coupled receptor, PtdIns(4,5)P_2_ is hydrolyzes into the second messengers, diacylglycerol and inositol 1,4,5-trisphosphate, followed by recycling of the components and resynthesis of PtdIns(4,5)P_2_ (Balla, 2013). Although phosphoinositide turnover occurs in the cytoplasmic leaflet, it is still unknown how PtdIns is asymmetrically distributed in the cytoplasmic leaflet. Here, we identified PtdIns flippases of the P4-ATPase family using synthesized NBD-PtdIns and highlight the importance of P4-ATPase activity for phosphoinositide turnover at the plasma membrane.

## Results and Discussion

### ATP10A can flip both NBD-PtdIns and NBD-PtdCho

To identify P4-ATPases capable of flipping PtdIns, we synthesized NBD-labeled PtdIns as previously described(Wang *et al*, 2018b) (Supplementary Figure S1). We first examined the flippase activities of plasma membrane-localized P4-ATPases which exhibit distinct substrate specificities; PtdSer flippases (ATP11A and ATP11C), PtdCho flippase (ATP10A), and glucosylceramide flippase (ATP10D) using NBD-labeled phospholipids (PtdIns, PtdCho, PtdSer, and SM)(Naito *et al*., 2015; Roland *et al*., 2019; Shin & Takatsu, 2022; Takatsu *et al*., 2014). Cells expressing ATP10A showed flippase activity toward both NBD-PtdIns and NBD-PtdCho, although the activity toward PtdIns was lower than that toward PtdCho (Figure 1A). By contrast, cells expressing PtdSer flippases (ATP11A and ATP11C) or glucosylceramide flippase (ATP10D) did not exhibit flippase activity toward PtdIns. Moreover, significant PtdIns flipping activity was not detected in cells expressing the ATPase-deficient mutant ATP10A(E203Q), demonstrating that ATP hydrolysis is essential for the PtdIns flipping activity (Figure 1B). The flippase activity of ATP10A toward NBD-PtdIns as well as NBD-PtdCho increased with time, whereas no flipping activity was observed toward NBD-PtdSer or NBD-SM (Figure 1B). These results indicate that ATP10A, initially identified as a PtdCho flippase, is also capable of translocating PtdIns. Contrary to previous reports that ATP8A1 can translocate PtdEtn at the plasma membrane (Kato *et al*, 2013) and PtdSer at endosomes (Lee *et al*, 2015), we could not detect flippase activities toward any examined phospholipids (including PtdEtn and PtdSer) or sphingolipids (Roland *et al*., 2019) in cells expressing ATP8A1 (Figure 1A).

**Figure 1.**
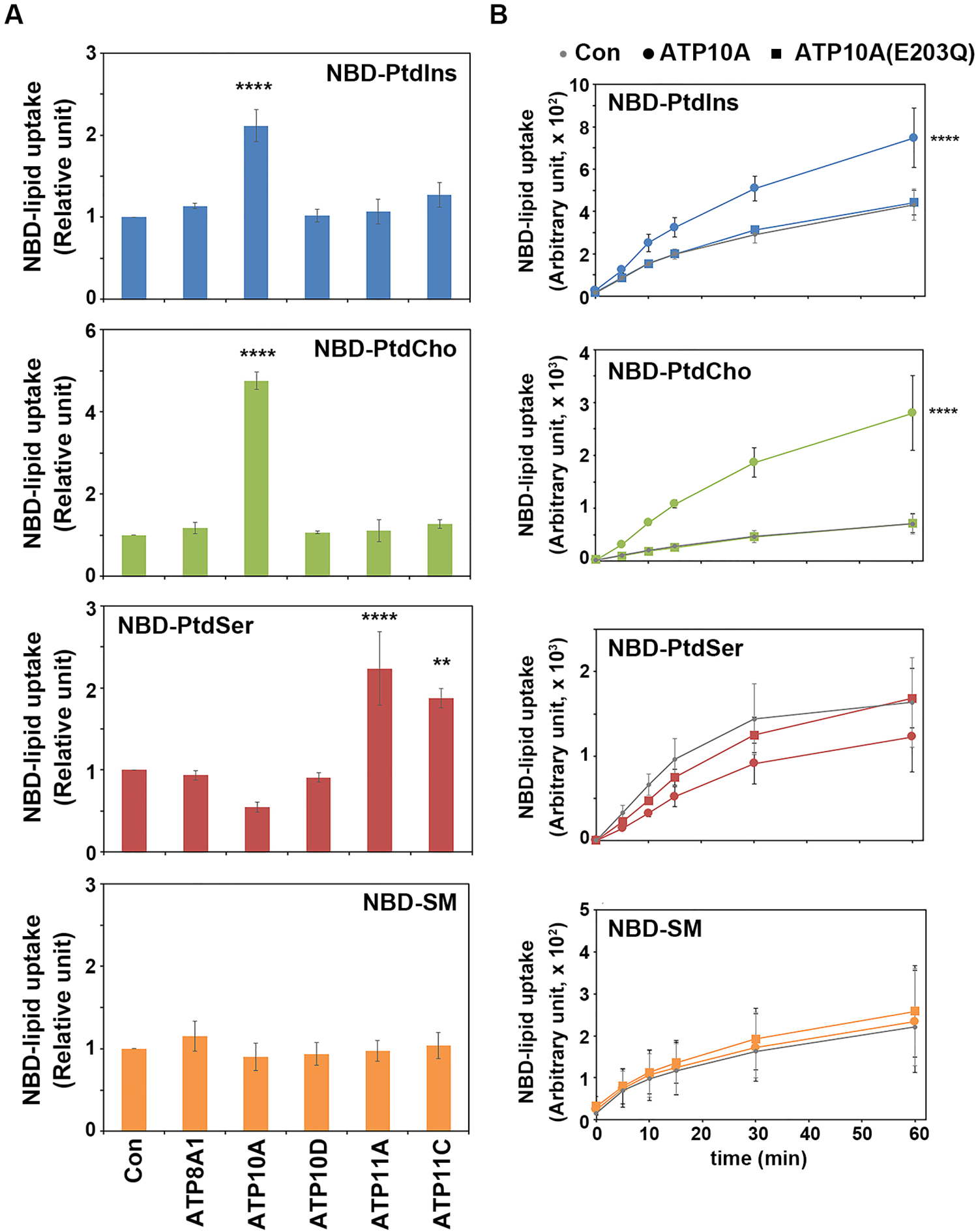
Flippase activities of P4-ATPases toward NBD-PtdIns. (**A**) HeLa cells stably expressing each P4-ATPase with C-terminal HA tag were incubated with the indicated NBD-phospholipids at 15° C for 15min (NBD-PtdIns, PtdCho, and SM) or 5 min (NBD-PtdSer). After extraction of labeled lipids in the exoplasmic leaflet of the cells with fatty acid-free BSA, the residual fluorescence intensity of the cells was measured by flow cytometry. Graphs are representative of at least three independent experiments, and results display averages of triplicates ± SD. Fold increase of NBD-lipid uptake compared to control cells (Con) is shown. A one-way analysis of variance (ANOVA) was performed to assess the variance, and comparisons with the control were performed using Dunnett analysis. (**B**) Control HeLa cells and cells stably expressing ATP10A-HA and ATP10A(E203Q)-HA, were incubated with the indicated NBD-lipids at 15° C for the indicated times (x axis). Graphs display averages of triplicates ± SD. A two-way ANOVA was performed to assess the variance, and comparisons with the control were performed using Dunnett analysis. ** p<0.01, *** p<0.001, **** p<0.0001.

### PtdCho flippases can flip PtdIns

Subsequently, we investigated whether the ability of other plasma membrane-localizing PtdCho flippases (ATP8B1 and ATP8B2) were able to translocate PtdIns. To this end, we established HeLa cells stably expressing ATP8B1, ATP8B2, and their ATPase-deficient mutants (EQ) and analyzed their flippase activity toward NBD-PtdIns, NBD-PtdCho, NBD-PtdSer, and NBD-SM. Notably, cells expressing ATP8B1 and ATP8B2 could also flip NBD-PtdIns. However, cells expressing the ATPase-deficient mutants ATP8B1(E234Q) or ATP8B2(E171Q), did not show any significant PtdIns flippase activity (Figure 2). Consistent with previous reports, cells expressing ATP8B1 of ATP8B2 did not display flippase activity towards NBD-PtdSer or NBD-SM (Figure 2) (Shin & Takatsu, 2022; Takatsu *et al*., 2014). These findings provide compelling evidence that PtdCho flippases can also translocate PtdIns. Additionally, we established HeLa cells stably expressing ATP8B4, a protein to closely related ATP8B1 and ATP8B2. However, ATP8B4 was unable to translocate any of the examined phospholipids and sphingolipids (Figure 2) (Roland *et al*., 2019; Takatsu *et al*., 2014) (Shin & Takatsu, 2022). Taken together, these results strongly support that the PtdCho flippases ATP8B1, ATP8B2, and ATP10A can efficiently translocate PtdIns in the plasma membrane.

**Figure 2.**
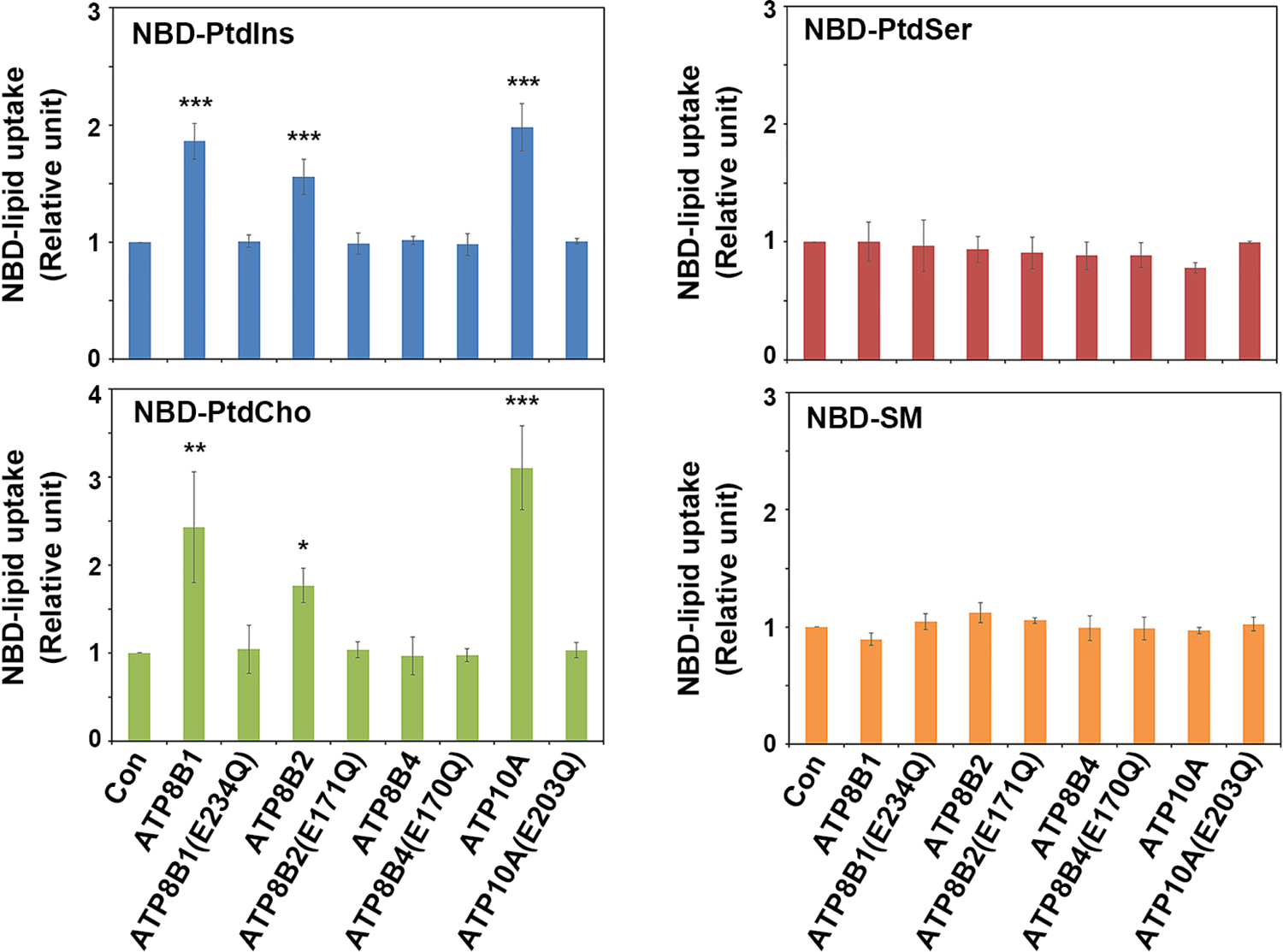
PtdIns-translocating activity of PtdCho flippases. HeLa cells stably expressing P4-ATPases with C-terminal HA tag, and their EQ mutants were incubated with the indicated NBD-phospholipids at 15° C for 15min (NBD-PtdIns, PtdCho, and SM) or 5 min (NBD-PtdSer). After extraction with fatty acid-free BSA, the residual fluorescence intensity of the cells was measured by flow cytometry. Graphs are representative of at least two independent experiments, and the data are averages of triplicates ± SD. The fold increase of NBD-lipid uptake compared with control cells (Con) is shown. A one-way ANOVA was performed to assess the variance, and comparisons with the control were performed using Dunnett analysis. * p<0.05, ** p<0.01, *** p<0.001.

### PtdIns(4,5)P_2_ rapidly disappears in CDC50A-KO cells upon Gq-coupled receptor activation

We then investigated the potential role of PtdIns flippases in phosphoinositide turnover. Since multiple P4-ATPases have been implicated in PtdIns flipping, we generated *CDC50A*-knockout (KO) cells using the CRISPR-Cas9 system (Supplementary Figure S2). CDC50A is essential for the transport of most P4-ATPases, including ATP8B1, ATP8B2, and ATP10A, from the endoplasmic reticulum to final destinations (Bryde *et al*., 2010; Naito *et al*., 2015; Takatsu *et al*., 2011; van der Velden *et al*., 2010b); it is of note that HeLa cells hardly express CDC50B (Bryde *et al*., 2010). Depletion of CDC50A in two independent KO clones was confirmed by immunoblotting (Supplementary Figure S2A). In addition, the decrease in PS flipping activity and the increase in PS exposure on the cell surface in *CDC50A*-KO cells were confirmed using NBD-PS and Annexin V staining, respectively (Supplementary Figure S2B and C), as previously reported for lymphocytes (Segawa *et al*, 2014). Disruption of all *CDC50A* alleles in the clone used in this study (clone 1-1) was confirmed by genomic sequencing (Supplementary Figure S2D).

To examine the subcellular distribution of phosphoinositides in both the parental HeLa and *CDC50A*-KO cells, we engineered these cells to stably express tagRFP (tRFP)-tagged phosphoinositide probes (Figure 3A). Specifically, tRFP-P4M was used for detection of PtdIns(4)P, the tRFP-PLCδ-PH domain for PtdIns(4,5)P_2_, and the tRFP-Akt-PH domain for PtdIns(3,4)P_2_ and PtdIns(3,4,5)P_3_ (Hammond *et al*, 2014; Shin *et al*, 2005; Stauffer *et al*, 1998). We observed that tRFP-P4M localized to the Golgi and the plasma membrane, and tRFP-PLCδ-PH and tRFP-Akt-PH domains mainly localized to the plasma membrane, as previously reported (Hammond *et al*., 2014; Shin *et al*., 2005; Stauffer *et al*., 1998). Importantly, we did not observe any significant differences in the distribution or intensity of the fluorescent signal between *CDC50A*-KO and control cells for any phosphoinositide probe. Furthermore, we examined the subcellular distribution and cellular levels of PtdIns(4)P and PtdIns(4,5)P_2_ at the steady state using specific antibodies (Figure 3B). Similarly, we did not observe any significant alterations in the distribution or levels of PtdIns(4)P or PtdIns(4,5)P_2_ between *CDC50A*-KO and parental control cells. These results suggest that, under steady state conditions, the total levels of each phosphoinositide remain largely unaltered in *CDC50A*-KO cells.

**Figure 3.**
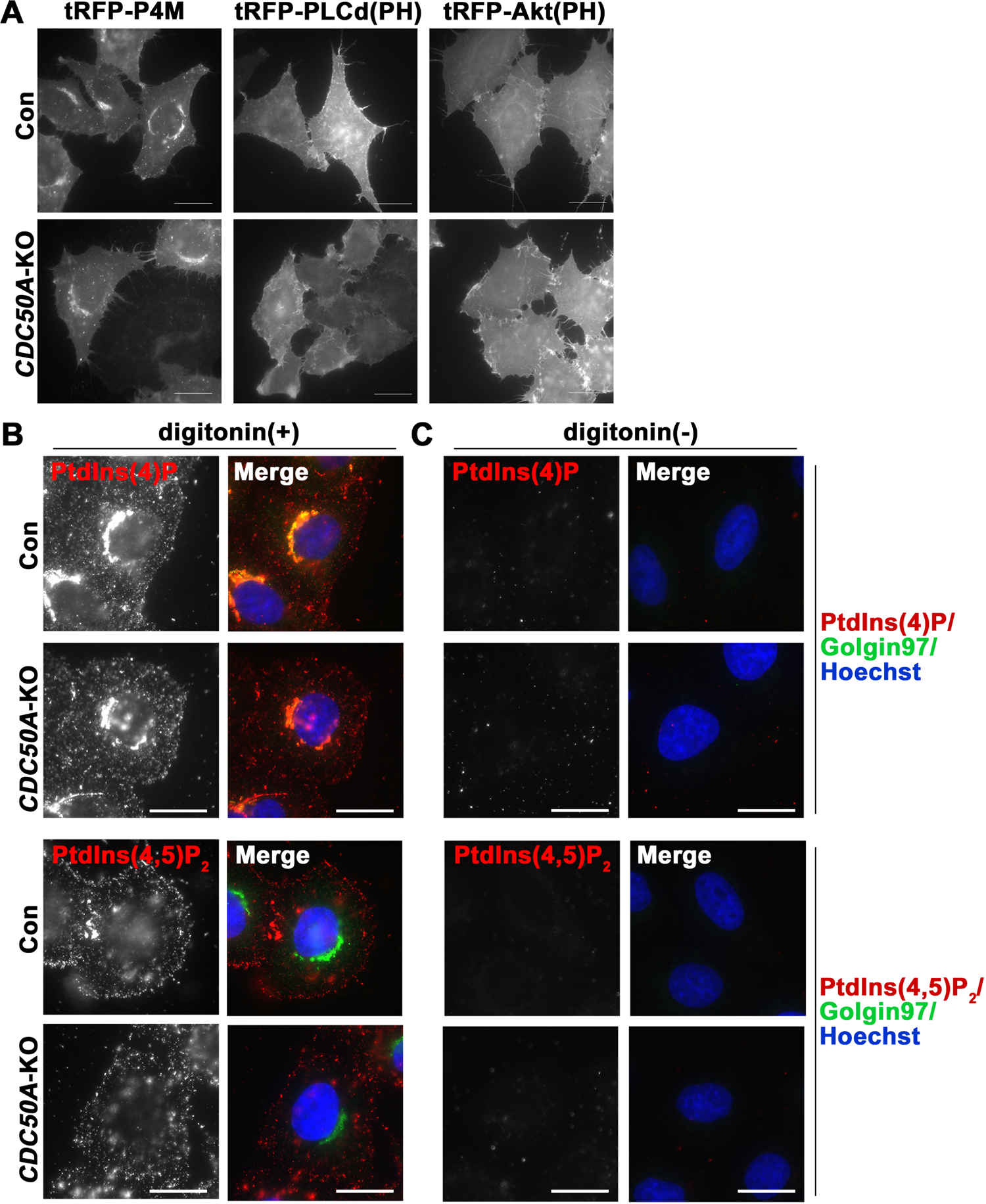
Distribution of phosphoinositides in *CDC50A*-KO cells. (**A**) Parental HeLa cells (Con) and *CDC50A*-KO cells stably expressing N-terminally tRFP-tagged P4M, PLCδ-PH domain, or Akt-PH domain for the detection of cellular PtdIns(4)P, PtdIns(4,5)P_2_, and PtdIns(3,4)P_2_/PtdIns(3,4,5)P_3_, respectively, were fixed, and the phosphoinositide distribution patterns were visualized using epifluorescence microscopy. (**B, C**) Parental HeLa and *CDC50A*-KO cells were fixed and either permeabilized with digitonin (**B**) or left non-permeabilized (**C**). The cells were subjected to immunostaining with anti-Golgin97 and either anti-PtdIns(4)P or anti-PtdIns(4,5)P_2_ antibody, followed by incubation with AlexaFluor488-conjugated anti-mouse IgG1 and anti-Cy3-conjugated anti-mouse IgM secondary antibodies. Hoechst33342 was added during the incubation with primary antibodies. Bars, 20 μm.

Since PtdSer are exposed at the cell surface in *CDC50A*-KO cells (Segawa *et al*., 2014) (Supplementary Figure S2C), PtdIns may exhibit similar behavior. However, there are currently no available antibodies that specifically recognize PtdIns. To examine whether PtdIns(4)P and PtdIns(4,5)P_2_ are exposed at the cell surface of *CDC50A*-KO cells, we performed immunofluorescence analysis, without permeabilizing the cells with digitonin, but were unable to detect these phosphoinositides at the cell surface (Figure 3C). This finding suggests that, in *CDC50A*-KO cells, phosphoinositides are unlikely to be exposed at the cell surface although we cannot rule out that PtdIns are exposed at the cell surface.

Next, we investigated the PtdIns(4,5)P_2_ turnover following the activation of Gq-coupled receptor. To this end, we established parental HeLa cells and *CDC50A*-KO cells stably expressing the Gq-coupled serotonin receptor (5-hydroxytryptamine [HT] receptor 2A; 5-HT_2A_). Upon activation of Gq-coupled receptors, PLC is activated, leading to the hydrolysis of PtdIns(4,5)P_2_ into inositol 1,4,5-trisphosphate and diacylglycerol (Figure 4A). We visualized the PtdIns(4,5)P_2_ levels using an anti-PtdIns4,5P_2_ antibody after treatment with serotonin (5-HT) in cells stably expressing 5-HT_2A_ (Figure 4B). Upon treatment with serotonin (5-HT), the PtdIns(4,5)P_2_ signals were rapidly decreased after 0.5 min, followed by a gradual recovery after 5 min (Figure 4B). Notably, the decrease in PtdIns(4,5)P_2_ signals was more pronounced in *CDC50A*-KO cells than in parental cells (Figure 4B, 0.5 and 2 min). The reduction and gradual recovery in the PtdIns(4,5)P_2_ signals was confirmed by quantification of the fluorescence intensity in individual cells using the ZEN software (Figure 4C). A significant reduction in the relative PtdIns(4,5)P_2_ signal intensity was observed at 0.5 min in *CDC50A*-KO cells compared with parental cells, although both parental and KO cells showed similar fluorescence intensities at 0 min (Figure 4D). These findings suggest that the available PtdIns(4,5)P_2_ levels, which can be rapidly hydrolyzed by PLC upon agonist activation, are lower in *CDC50A*-KO cells than in control cells. This decrease could be attributed to the absence of PtdIns flippases at the plasma membrane. However, the diminished PtdIns(4,5)P_2_ levels at the plasma membrane were not detected at steady state (Figures 3B, 4B, and C). Indeed, the PtdIns(4,5)P_2_ levels can be restored by PtdIns transfer proteins, which transport PtdIns from the ER upon agonist stimulation (Chang *et al*, 2013; Chang & Liou, 2015; Kauffmann-Zeh *et al*, 1995; Kim *et al*, 2015).

**Figure 4.**
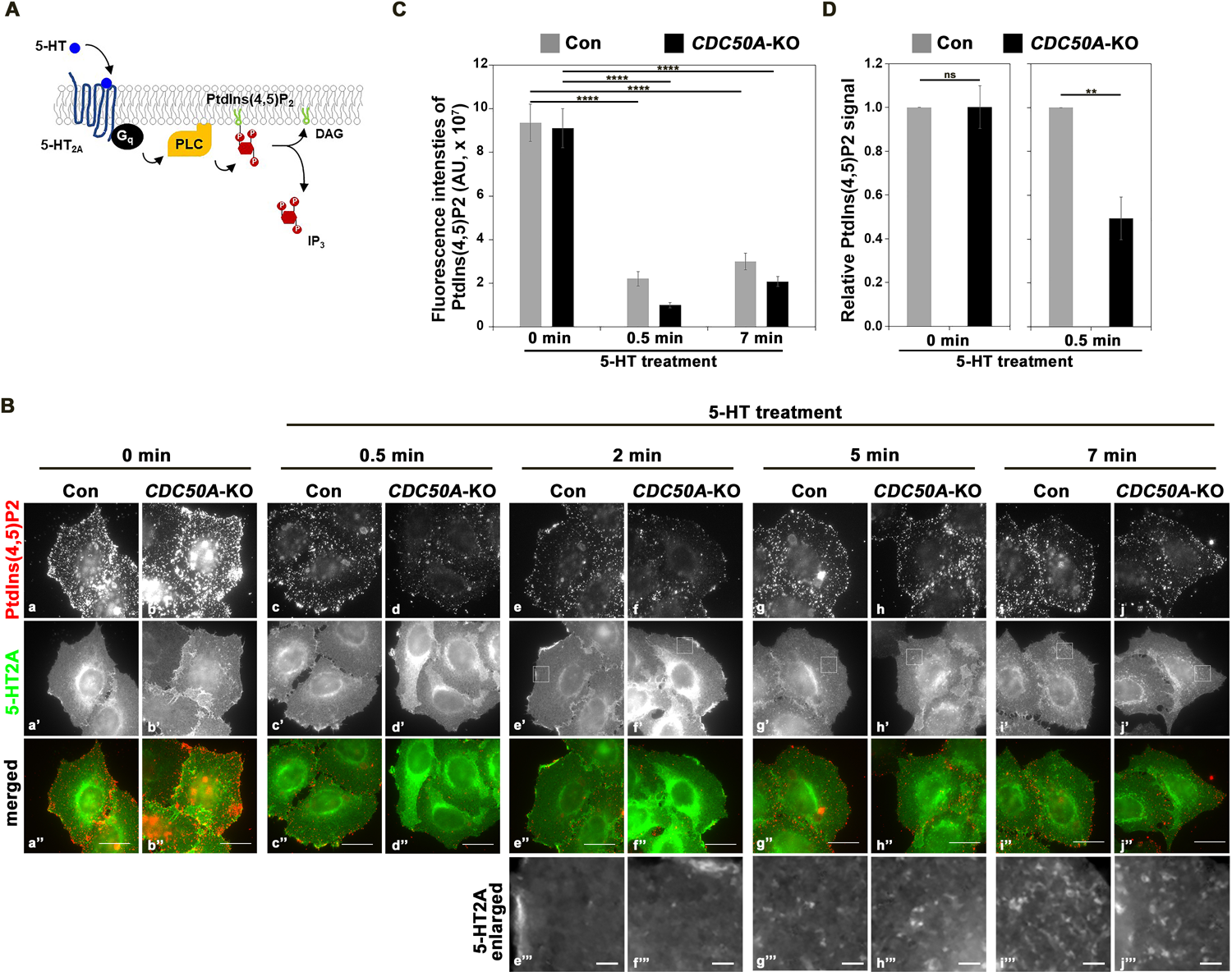
PtdIns(4,5)P_2_ turnover in *CDC50A*-KO cells. (**A**) Schematic representation of PtdIns(4,5)P_2_ hydrolysis upon 5-HT stimulation. (**B**) Parental HeLa (Con) and *CDC50A*-KO cells stably expressing C-terminally FLAG-tagged 5-HT_2A_ were serum-starved for 3 h and then treated with 1 μM 5-HT for the indicated times. The cells were fixed and stained with anti-DYKDDDDK and anti-PtdIns(4,5)P_2_ antibodies, followed by incubation with AlexaFluor488-conjugated anti-mouse IgG2b and Cy3-conjugated anti-mouse IgM secondary antibodies. Hoechst33342 was added during the incubation with primary antibodies. Bars, 20 μm, and 2 μm in enlarged images. (**C, D**) The fluorescence intensity of the PtdIns(4,5)P_2_ signal was measured using the ZEN software. (**C**) Graphs display the average ± SEM of representative data from more than two independent experiments. Approximately 40 cells in each sample were analyzed. A one-way ANOVA was performed to assess the variance, and comparisons to 0 min were performed using Tukey’s post-hoc analysis. (**D**) Graphs display the average ± SD from three independent experiments. Over 30 cells were analyzed in each sample, and the fold changes compared to the control at each time point are presented. Student’s t tests were performed for pairwise comparisons. **** p<0.0001, ** p<0.01, ns, not significant.

Upon treatment with 5-HT, the 5-HT_2A_ was endocytosed (Takatsu *et al*, 2017) and appeared in endosomal puncta after 5 min (Figure 4Bg’’’ and h’’’). The endosomal signal became more prominent after 7 min (Figure 4Bi’’’ and j’’’). These observations indicate a correlation between the sequestration of the 5-HT_2A_ and the recovery of the PtdIns(4,5)P_2_ fluorescent signals.

Mutations in the human *ATP8B1* gene cause PFIC1 (Folmer *et al*., 2009; Paulusma *et al*., 2006). Some variants of PFIC1 ATP8B1 are unable to flip PC (Takatsu *et al*., 2014) and others fail to express due to defects in protein folding and subsequent degradation (Takatsu *et al*., 2014; van der Velden *et al*, 2010a). In addition to the loss of PtdCho flippase activity (Shin & Takatsu, 2019), the absence of PtdIns flippase activity may also contribute to the development of PFIC1. Phosphoinositide 3-kinase activity is involved in the endocytosis of bile salt export pump (BSEP), encoded by the gene responsible for PFIC2, and in the secretion of bile salt (Boaglio *et al*, 2010). Therefore, the PtdIns flippase activity of ATP8B1 may influence the function of BSEP and/or intracellular trafficking of BSEP via the phosphoinositide 3-kinase signaling cascade. The responsible gene for PFIC3 is *ABCB4*, which acts as a PtdCho floppase by translocating PtdCho from inner to the outer leaflet of the plasma membrane. It is worth noting that similar clinical manifestations are observed in PFIC1 and PFIC2 compared to PFIC3 patients; 1) both PFIC1 and PFIC2 exhibit low concentration of primary bile acids within the bile composition, while PFIC3 shows low phospholipid concentration, 2) PFIC1 and PFIC2 usually manifest in the first months of life, whereas onset of PFIC3 occurs later, 3) severe pruritus is usually observed in PFIC1 and PFIC2 patients (Davit-Spraul *et al*, 2009; Jacquemin, 2012).

The exogenous expression of ATP10A in HeLa cells induces the formation of inward membrane curvature (Takada *et al*., 2018) and cell shape changes (Miyano *et al*., 2016; Naito *et al*., 2015). These phenotypes potentially are attributed to the alteration in the transbilayer balance of phospholipid mass. Considering that more than 90 % of PtdCho is estimated to be present in the outer leaflet of the HeLa plasma membrane (Murate *et al*., 2015), these phenotypes are likely associated with the enhanced PtdCho-flipping activity resulting from the exogenous expression of ATP10A. Additionally, the ATP10A expression facilitates endocytosis (Takada *et al*, 2018) and delays cell adhesion and spreading on the extracellular matrix (Miyano *et al*, 2016; Naito *et al*., 2015). The enhanced PtdIns-flipping activity, resulting from the exogenous expression of ATP10A, may contribute to these phenotypes since phosphoinositides play critical roles in endocytosis and actin cytoskeleton rearrangement (Balla, 2013; Nebl *et al*, 2000; Posor *et al*., 2022; Schink *et al*., 2016).

The mouse *Atp10a* gene is linked to type-2 diabetes phenotypes (Dhar *et al*., 2004) and certain single-nucleotide polymorphisms in ATP10A are associated with insulin resistance (Irvin *et al*., 2011). Regulation of phosphoinositides, particularly PtdIns(3,4,5)P_3_, at the plasma membrane is crucial for insulin signaling mediated by phosphoinositide 3-kinase and Akt cascade (Saltiel, 2021) (J.P.Kirwan & Aguila, 2003). Therefore, a defect in PtdIns-flippase activity of ATP10A may contribute to impaired insulin signaling and the development of type-2 diabetes.

While it remains unclear whether PtdIns flippases (ATP8B1, ATP8B2, and ATP10A) are directly involved in the rapid turnover of PtdIns(4,5)P_2_ at the plasma membrane in response to agonist stimulation, our findings provide novel insight into the phosphoinositide signaling pathway. During the preparation of this manuscript, a preprint was published reporting that the ATPase activity of ATP8B1 can be stimulated by various phospholipids including PtdIns, which supports our observations (Thibaud *et al*, 2023). To further advance our understanding of the physiological roles of PtdIns-flipping in cells, it will be essential to identify the specific amino acids responsible for differences in substrate specificity between PtdIns and PtdCho.

## Materials and Methods

### Plasmids

Partial cDNA fragments of ATP8B2 were obtained from Kazusa (ORK04226) and the 5’ fragments of ATP8B2 were obtained by RT-PCR from total RNA of HeLa cells using the following primer pairs: sense, 5’-ggcgtcgacaccatggcagtgtgtgcaaaaaagcgc-3’, antisense, 5’-ccagaggattgaaggagaagtc-3’). The complete human ATP8B2 cDNA (NM_001370597) was inserted into pENTR3C (Thermo Fisher Scientific, Waltham, MA, USA) and transferred to expression vectors using the Gateway system (Thermo Fisher Scientific) as previously described (Takatsu *et al*., 2011; Takatsu *et al*., 2014). Point mutations were introduced into ATP8B2 and ATP8B4 using the QuikChange II XL site-directed mutagenesis kit (Agilent Technologies, Santa Clara, CA, USA) and/or the SLiCE cloning method (Zhang *et al*, 2012). The expression vector for the mouse 5-HT2A receptor was described previously (Takatsu *et al*., 2017). The cDNAs of P4M (plasmid #51472) (Hammond *et al*., 2014) and PLCδ-PH domain (plasmid #21179) (Stauffer *et al*., 1998) were obtained from Addgene. The tRFP cDNA was a gift from Hideki Shibata (Nagoya University) (Shibata *et al*, 2010), and the Akt-PH domain cDNA was a gift from Marino Zerial (MPI-CBG). To generate the expression vector for the tRFP-tagged P4M and PLCd-PH domain, the cDNAs were inserted into the pRRLsinPPT-tRFP and the pMXs-tagRFP-puro vectors, respectively. The blasticidin-resistant gene was subsequently substituted with the puromycin-resistant gene. The Akt-PH domain cDNA was inserted into the pMXs-tagRFP-puro vector.

### Antibodies and reagents

Anti-CDC50A antisera were raised in rabbits as described previously (Tone *et al*, 2020). Monoclonal mouse anti-PtdIns(4)P and anti-PtdIns(4,5)P_2_ IgM antibodies were purchased from Echelon Biosciences (Salt Lake City, UT, USA), anti-DYKDDDDK (1E6) was purchased from Wako Pure Chemical Industries (Osaka, Japan), anti-Golgin97 antibody (CDF4) was purchased from Thermo Fisher Scientific, and anti-β-actin antibody (C4) was purchased from Santa Cruz (Dallas, TX, USA). AlexaFluor-conjugated secondary antibodies were purchased from Thermo Fisher Scientific. Cy3- and horseradish peroxidase-conjugated secondary antibodies were purchased from Jackson ImmunoResearch Laboratories (West Grove, PA, USA). NBD-PtdSer (1-oleoyl-2-[6-[(7-nitro-2-,3-benzoxadiazol-4-yl)amino]hexanoyl]-*sn*-glycero-3-phosphoserine), NBD-PtdEtn (1-oleoyl-2-[6-[(7-nitro-2-,3-benzoxadiazol-4-yl)amino]hexanoyl]-*sn*-glycero-3-phosphoethanolamine), NBD-PtdCho (1-oleoyl-2-[6-[(7-nitro-2-,3-benzoxadiazol-4-yl)amino]hexanoyl]-*sn*-glycero-3-phosphocholine), and NBD-SM (*N*-[6-[(7-nitro-2-,3-benzoxadiazol-4-yl)amino]hexanoyl]-sphingosine 1-phosphocholine) were purchased from Avanti Polar Lipids (Ablabaster, AL, USA). 5-hydroxytryptamine and propidium iodide (PI) were purchased from Nacalai Tesque (Kyoto, Japan). FITC-Annexin V was purchased from BioLegend (San Diego, CA, USA). Hoechst33342 was purchased from Thermo Fisher Scientific.

### Cell culture and establishment of stable cells

HeLa cells were cultured in minimum essential medium (MEM, Nacalai Tesque), supplemented with MEM non-essential amino acids (Nacalai Tesque) and 10% fetal calf serum (Gibco, Waltham, MA, USA). HeLa cells stably expressing C-terminally HA-tagged P4-ATPases were described previously (Naito *et al*., 2015; Roland *et al*., 2019; Takatsu *et al*., 2014). HeLa cells stably expressing C-terminally HA-tagged ATP8B2, ATP8B2(E171Q) and ATP8B4(E170Q) were established as previously described (Takatsu *et al*., 2014). For the generation of HeLa and *CDC50A*-KO cells stably expressing each phosphoinositide probe, recombinant retrovirus for the expression of tRFP-PLCδ(PH) and tRFP-Akt(PH) and recombinant lentivirus for the expression of tRFP-P4M were produced and used to infect HeLa and *CDC50A*-KO cells. To generate stable cells expressing tRFP-PLCδ(PH) were selected in medium containing blasticidin (10μg/ml) and a pool of cells expressing tRFP-P4M was isolated using an SH800S cell sorter (SONY biotechnology, San Jose, CA, USA). Although drug selection was not used to isolate cells stably expressing tRFP-Akt(PH), almost 100% of the cells were found to express tRFP-Akt(PH). To establish HeLa and *CDC50A*-KO cells that stably express C-terminally FLAG-tagged 5-HT2A-R, recombinant retrovirus for expression of 5-HT2A-R-FLAG was produced and used to infect the cells (Takatsu *et al*., 2017). Although the infected cells were not drug-selected, over 80% of the cells expressed 5-HT2A-R-FLAG stably. For 5-HT treatment, cells on coverslips were incubated with MEM supplemented 2% BSA for 3 h, followed by incubation with 1 μM of 5-HT in MEM, supplemented with 2% BSA and 25 mM HEPES-KOH pH 7.4 for the indicated times. The cells were then immediately fixed and subjected to immunofluorescence analysis.

### Gene editing using the CRISPR/Cas9 system

The *CDC50A* gene was disrupted in HeLa cells using the CRISPR/Cas9 system. Target sequences for the *CDC50A* gene were designed using the CRISPR Design Tool from the Zhang lab (http://crispr.mit.edu). Complementary oligonucleotides (#1, 5′-caccg<u>actcggagaccggataaca</u>-3′ and 5′-aaac<u>tgttatccggtctccgagtc</u>-3′, #2, 5’-cacc<u>gcggtgccccccggagcaca</u>-3’ and 5’-aaac<u>tgtgctccggggggcaccgc-</u>3’; target sequences are underlined) were synthesized and introduced into the BbsI-digested vector PX459 (Addgene plasmid #48139). To edit the *CDC50A* gene, we used a previously described method (Nozaki *et al*, 2017; Tanaka *et al*, 2016), which involved the co-transfection of the plasmid containing the *CDC50A* target sequences and the *Cas9* gene with a donor plasmid (pDonor-tBFP-NLS-Neo; Addgene plasmid #80766) referred to as Donor. Both plasmids were transfected into HeLa cells using the X-tremeGENE9 DNA Transfection Reagent (Roche) and selected in medium containing G418 (1 mg/mL). Cell clones were isolated using the SH800S cell sorter. The KO was confirmed by immunoblot analysis and the flippase assay using NBD-PtdSer. Genomic DNA was extracted from clone 1-1 (used for further experiments in this study), and PCR was performed to amplify the region of interest. The KO was confirmed by direct sequencing of the amplified PCR product, which revealed the presence of mutations in three alleles (Supplementary Figure S2).

### Immunofluorescence analysis

Immunostaining was performed as previously described (Shin *et al*, 2004), with some modifications. Cells were fixed with 3% PFA for 20 min at RT and quenched with 50 mM NH_4_Cl for 5 min at RT. They were then permeabilized with 20 μM digitonin for 5 min at RT and blocked with 10% FCS for 15 min. Cells were incubated with primary antibodies followed by secondary antibodies and mounted on a slide glass with Mowiol as previously described (Shin *et al*., 2004). Samples were visualized using an Axiovert 200MAT microscope (Carl Zeiss, Jena, Germany) and the fluorescence intensities of PtdIns4,5P_2_ were analyzed using the ZEN software (Carl Zeiss).

### NBD-labeled PtdIns synthesis

NBD-PtdIns (1-oleoyl-2-[6-[(7-nitro-2-,3-benzoxadiazol-4-yl)amino]hexanoyl]-*sn*-glycero-3-phosphoinositol) was synthesized from NBD-PtdCho and *myo*-inositol by phospholipase D (PLD)-catalyzed transphosphatidylation, as previously reported (Wang *et al*., 2018b; Wang *et al*, 2022). A microbial-PLD engineered variant bearing three amino acid replacements (G186T/W187N/Y385R) (Damnjanovic *et al*, 2016) was used as the catalyst. One milligram of C18:1/NBD-C6-PtdCho (Avanti) dissolved in 100 µL of ethyl acetate was mixed with 80 µL of NaCl-saturated 50 mM acetate buffer (pH 5.6) containing 18 mg of *myo*-inositol and 20 µL of 0.5 mg/mL PLD variant. The mixture was incubated at 20° C with vigorous shaking for 24 h. Ten microliters of 1 M HCl were added to stop the reaction, and the lipid was extracted with 200 µL of chloroform/methanol (2:1). From the resultant extract, NBD-PtdIns was purified by silica gel column chromatography. The purified product exhibited a single spot in thin layer chromatography (TLC) analysis with a trace of non-polar fluorescent impurities (Supplementary Figure S1A). The product displayed an *m/z* value of 873 (M-H)^-^ on LC-MS analysis (theoretical C_39_H_63_N_4_O_16_P = 874.4) (Supplementary Figure S1B).

### Cell-based flippase assay

Incorporation of NBD-lipids was analyzed by flow cytometry as described previously (Takatsu *et al*., 2014). In brief, HeLa cells were detached from dishes in PBS containing 5 mM EDTA and harvested by centrifugation. Cells (2 × 10^5^ cells/sample) were washed and equilibrated at 15° C for 15 min in 100 μL Hank’s balanced salt solution (pH 7.4) containing 1 g/L glucose (HBSS-glucose). An equal volume of 2 μM NBD-phospholipid in HBSS-glucose was added to the cell suspension, which was further incubated at 15° C for 5 or 15 min. Two hundred microliters of cell suspension were mixed with 200 μL ice-cold HBSS-glucose containing 5% fatty acid-free BSA (Wako) to extract NBD-lipids incorporated into the exoplasmic leaflet of the plasma membrane, as well as unincorporated lipids. Subsequently, the cells were analyzed by flow cytometry using FACS Calibur (BD Biosciences) to measure the fluorescence of incorporated NBD-lipids and lipids translocated into the cytoplasmic leaflet of the plasma membrane. Data were analyzed with the FlowJo software (BD Biosciences).

### Annexin V assay

To detect PtdSer on the cell surface, HeLa cells were detached from dishes in PBS containing 2.5 mg/ml Trypsin, HBSS-glucose was added, and the cells were harvested by centrifugation. Cells were washed twice with ice-cold Annexin buffer (10 mM HEPES-KOH, pH 7.4, 140 mM NaCl, and 2.5 mM CaCl_2_) and suspended in ice-cold Annexin buffer. Then the same volume of FITC-Annexin V in ice-cold Annexin buffer was added, and the cells were incubated for 15 min on ice. Propidium iodide was added, and the cells were further incubated for 3 min on ice, harvested by centrifugation, resuspended in Annexin buffer, and then analyzed by flow cytometry using FACS Calibur. Data were analyzed using the FlowJo software.

## Acknowledgments

We thank Toshio Kitamura (University of Tokyo) and Hiroyuki Miyoshi (RIKEN BioResource Center) for providing plasmids for retroviral infection, Peter McPherson (McGill University) for providing plasmids for lentiviral infection, Marino Zerial (MPI-CBG) for the Akt-PH plasmid, and Hideki Shibata (Nagoya University) for the tRFP plasmid. This work was supported by JSPS KAKENHI Grant Numbers JP23H02434 (to H.-W.S.), JP20H03209 (to H.-W.S.), and JP20K07325 (to H.T.), the Takeda Science Foundation (to H.-W.S.), the Mizutani Glycoscience Foundation (to H.-W.S.), and the ONO Medical Research Foundation (to H.-W.S.).

## Supplementary Figure Legends

**Supplementary Figure S1.**
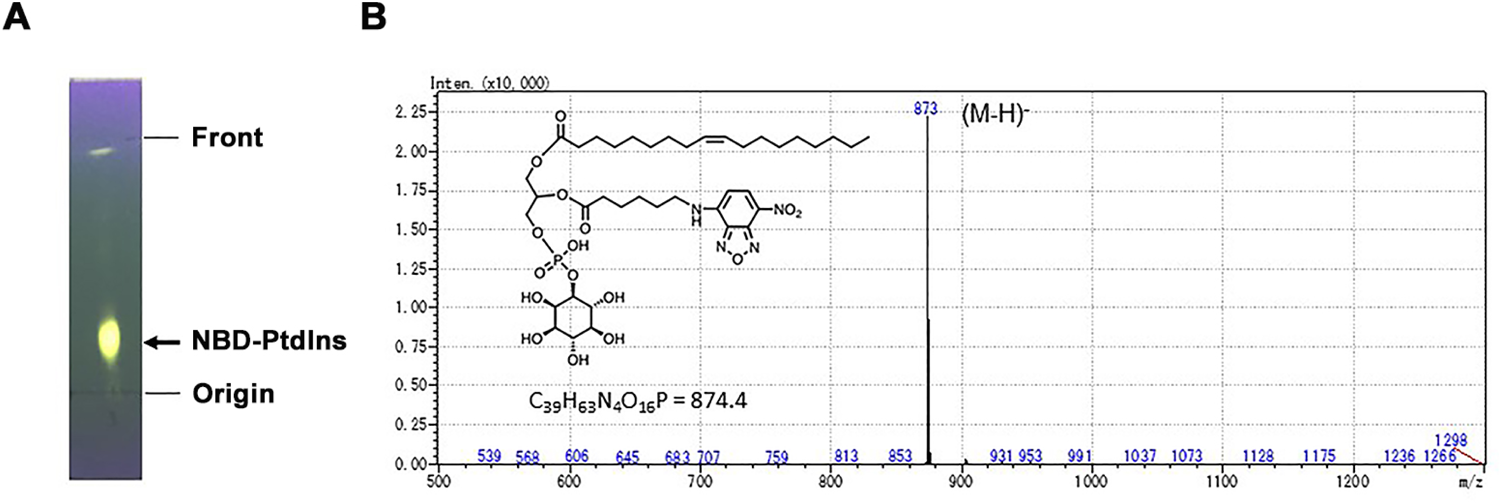
Preparation of NBD-PtdIns. (**A**) TLC analysis of the prepared NBD-PtdIns. TLC was run using chloroform/petroleum ether/methanol/acetic acid (4:3:2:1) as the developing solvent, and spots were visualized under UV light (365 nm). (**B**) Mass spectrum of the purified NBD-PtdIns. The purified lipid was analyzed by electrospray ionization mass spectrometry in negative-ion mode.

**Supplementary Figure S2.**
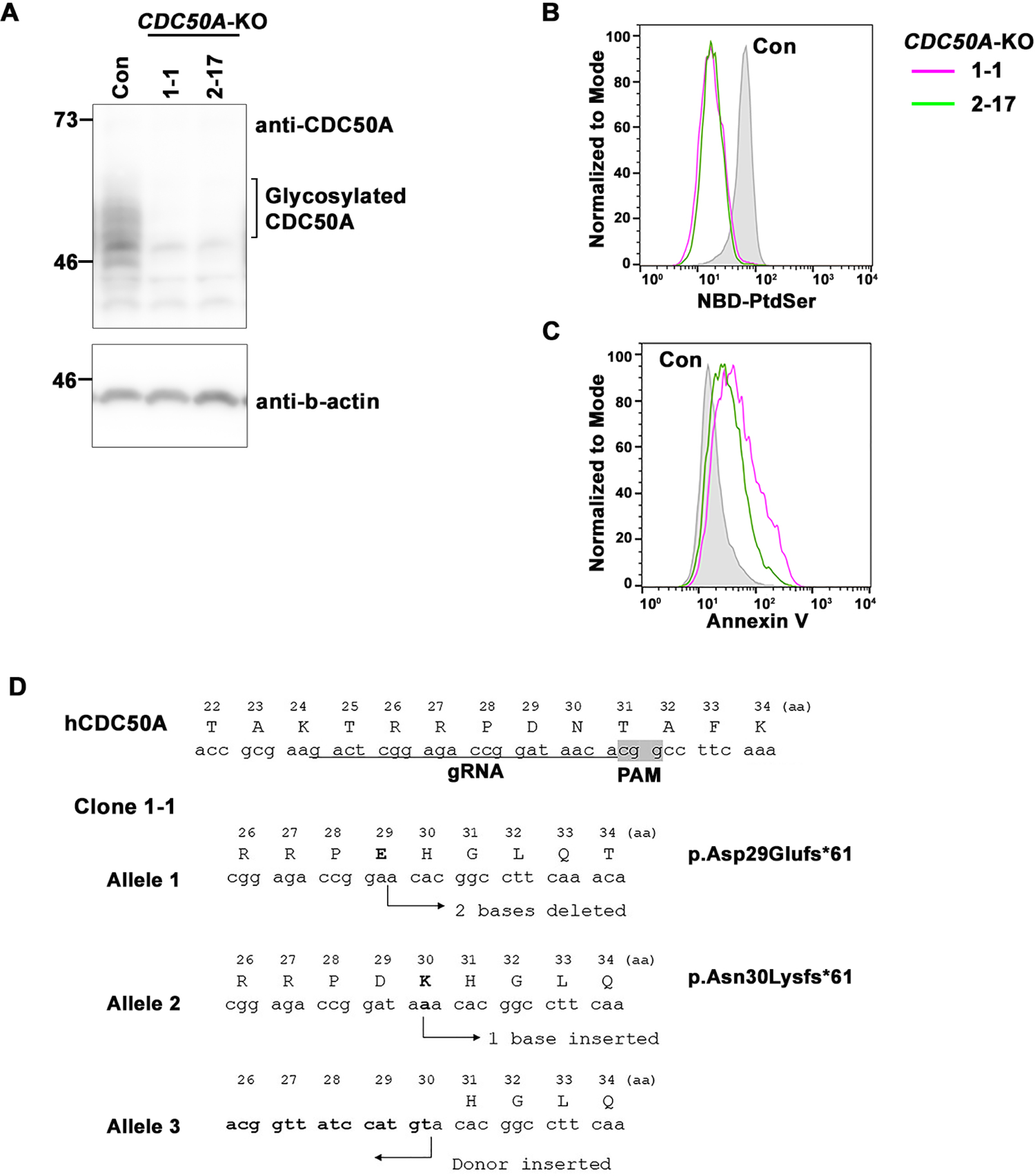
Disruption of *CDC50A* by CRISPR/Cas9 in HeLa cells. (**A**) Cell lysates were prepared from two clones of *CDC50A*-KO cells and subjected to immunoblot analysis using affinity-purified anti-CDC50A antibody to confirm the depletion of CDC50A proteins. Clone 1-1 and clone 2-17 were generated using two sgRNAs, #1 and #2, respectively. (**B**) Depletion of CDC50A was confirmed by flippase activity assay using NBD-PtdSer. Both independent clones exhibited a significant reduction in the uptake of NBD-PtdSer. (**C**) Depletion of CDC50A was confirmed by Annexin V assay. Both clones exhibited a significant increase in PtdSer exposure at the cell surface. (**D**) Genomic DNA analysis for clone 1-1 used in this study. *CDC50A* carries two independent indels and the Donor, indicating the presence of three *CDC50A* copies in the genome.

## References

Andersen JP, Vestergaard AL, Mikkelsen SA, Mogensen LS, Chalat M, Molday RS (2016) P4-ATPases as Phospholipid Flippases-Structure, Function, and Enigmas. Front Physiol 7: 275

Balla T (2013) Phosphoinositides: tiny lipids with giant impact on cell regulation. Physiological reviews 93: 1019–1137

Boaglio AC, Zucchetti AE, Sanchez Pozzi EJ, Pellegrino JM, Ochoa JE, Mottino AD, Vore M, Crocenzi FA, Roma MG (2010) Phosphoinositide 3-kinase/protein kinase B signaling pathway is involved in estradiol 17beta-D-glucuronide-induced cholestasis: complementarity with classical protein kinase C. Hepatology 52: 1465–1476

Bryde S, Hennrich H, Verhulst PM, Devaux PF, Lenoir G, Holthuis JC (2010) CDC50 Proteins Are Critical Components of the Human Class-1 P4-ATPase Transport Machinery. J Biol Chem 285: 40562–40572

Chang CL, Hsieh TS, Yang TT, Rothberg KG, Azizoglu DB, Volk E, Liao JC, Liou J (2013) Feedback regulation of receptor-induced Ca2+ signaling mediated by E-Syt1 and Nir2 at endoplasmic reticulum-plasma membrane junctions. Cell Rep 5: 813–825

Chang CL, Liou J (2015) Phosphatidylinositol 4,5-Bisphosphate Homeostasis Regulated by Nir2 and Nir3 Proteins at Endoplasmic Reticulum-Plasma Membrane Junctions. J Biol Chem 290: 14289–14301

Damnjanovic J, Kuroiwa C, Tanaka H, Ishida K, Nakano H, Iwasaki Y (2016) Directing positional specificity in enzymatic synthesis of bioactive 1-phosphatidylinositol by protein engineering of a phospholipase D. Biotechnol Bioeng 113: 62–71

Davit-Spraul A, Gonzales E, Baussan C, Jacquemin E (2009) Progressive familial intrahepatic cholestasis. Orphanet J Rare Dis 4: 1

Devaux PF (1991) Static and dynamic lipid asymmetry in cell membranes. Biochemistry 30: 1163–1173

Dhar MS, Sommardahl CS, Kirkland T, Nelson S, Donnell R, Johnson DK, Castellani LW (2004) Mice heterozygous for Atp10c, a putative amphipath, represent a novel model of obesity and type 2 diabetes. J Nutr 134: 799–805

Folmer DE, Elferink RPJO, Paulusma CC (2009) P4 ATPases - Lipid flippases and their role in disease. Biochimica et Biophysica Acta (BBA) - Molecular and Cell Biology of Lipids 1791: 628–635

Hammond GR, Machner MP, Balla T (2014) A novel probe for phosphatidylinositol 4-phosphate reveals multiple pools beyond the Golgi. J Cell Biol 205: 113–126

Hokin MR, Hokin LE (1953) Enzyme secretion and the incorporation of P32 into phospholipides of pancreas slices. J Biol Chem 203: 967–977

Irvin MR, Wineinger NE, Rice TK, Pajewski NM, Kabagambe EK, Gu CC, Pankow J, North KE, Wilk JB, Freedman BI et al (2011) Genome-wide detection of allele specific copy number variation associated with insulin resistance in African Americans from the HyperGEN study. PLoS One 6: e24052

J.P.Kirwan, Aguila LFd (2003) Insulin signalling, exercise and cellular integrity. Biochemical society transactions 31: 1281–1285

Jacquemin E (2012) Progressive familial intrahepatic cholestasis. Clinics and Research in Hepatology and Gastroenterology 36, Supplement 1: S26–S35

Kato U, Inadome H, Yamamoto M, Emoto K, Kobayashi T, Umeda M (2013) Role for Phospholipid Flippase Complex of ATP8A1 and CDC50A Proteins in Cell Migration. Journal of Biological Chemistry 288: 4922–4934

Kauffmann-Zeh A, Thomas GM, Ball A, Prosser S, Cunningham E, Cockcroft S, Hsuan JJ (1995) Requirement for phosphatidylinositol transfer protein in epidermal growth factor signaling. Science 268: 1188–1190

Kim YJ, Guzman-Hernandez ML, Wisniewski E, Balla T (2015) Phosphatidylinositol-Phosphatidic Acid Exchange by Nir2 at ER-PM Contact Sites Maintains Phosphoinositide Signaling Competence. Dev Cell 33: 549–561

Lee S, Uchida Y, Wang J, Matsudaira T, Nakagawa T, Kishimoto T, Mukai K, Inaba T, Kobayashi T, Molday RS et al (2015) Transport through recycling endosomes requires EHD1 recruitment by a phosphatidylserine translocase. EMBO J 34: 669–688

Lopez-Marques RL, Gourdon P, Gunther Pomorski T, Palmgren M (2020) The transport mechanism of P4 ATPase lipid flippases. The Biochemical journal 477: 3769–3790

Miyano R, Matsumoto T, Takatsu H, Nakayama K, Shin HW (2016) Alteration of transbilayer phospholipid compositions is involved in cell adhesion, cell spreading, and focal adhesion formation. FEBS Lett 590: 2138–2145

Murate M, Abe M, Kasahara K, Iwabuchi K, Umeda M, Kobayashi T (2015) Transbilayer distribution of lipids at nano scale. J Cell Sci 128: 1627–1638

Naito T, Takatsu H, Miyano R, Takada N, Nakayama K, Shin HW (2015) Phospholipid Flippase ATP10A Translocates Phosphatidylcholine and Is Involved in Plasma Membrane Dynamics. J Biol Chem 290: 15004–15017

Nebl T, Oh SW, Luna EJ (2000) Membrane cytoskeleton: PIP(2) pulls the strings. Current biology: CB 10: R351–354

Nozaki S, Katoh Y, Terada M, Michisaka S, Funabashi T, Takahashi S, Kontani K, Nakayama K (2017) Regulation of ciliary retrograde protein trafficking by the Joubert syndrome proteins ARL13B and INPP5E. J Cell Sci 130: 563–576

Paulusma CC, Groen A, Kunne C, Ho-Mok KS, Spijkerboer AL, Rudi de Waart D, Hoek FJ, Vreeling H, Hoeben KA, van Marle J et al (2006) Atp8b1 deficiency in mice reduces resistance of the canalicular membrane to hydrophobic bile salts and impairs bile salt transport. Hepatology 44: 195–204

Posor Y, Jang W, Haucke V (2022) Phosphoinositides as membrane organizers. Nat Rev Mol Cell Biol

Roland BP, Naito T, Best JT, Arnaiz-Yepez C, Takatsu H, Yu RJ, Shin HW, Graham TR (2019) Yeast and human P4-ATPases transport glycosphingolipids using conserved structural motifs. J Biol Chem 294: 1794–1806

Saltiel AR (2021) Insulin signaling in helath and disease. The journal of clinical investigation 131: e141141

Schink KO, Tan KW, Stenmark H (2016) Phosphoinositides in Control of Membrane Dynamics. Annual review of cell and developmental biology 32: 143–171

Segawa K, Kurata S, Yanagihashi Y, Brummelkamp TR, Matsuda F, Nagata S (2014) Caspase-mediated cleavage of phospholipid flippase for apoptotic phosphatidylserine exposure. Science 344: 1164–1168

Shibata H, Inuzuka T, Yoshida H, Sugiura H, Wada I, Maki M (2010) The ALG-2 binding site in Sec31A influences the retention kinetics of Sec31A at the endoplasmic reticulum exit sites as revealed by live-cell time-lapse imaging. Bioscience, biotechnology, and biochemistry 74: 1819–1826

Shin H-W, Hayashi M, Christoforidis S, Lacas-Gervais S, Hoepfner S, Wenk MR, Modregger J, Uttenweiler-Joseph S, Wilm M, Nystuen A et al (2005) An enzymatic cascade of Rab5 effectors regulates phosphoinositide turnover in the endocytic pathway. J Cell Biol 170: 607–618

Shin HW, Morinaga N, Noda M, Nakayama K (2004) BIG2, a guanine nucleotide exchange factor for ADP-ribosylation factors: its localization to recycling endosomes and implication in the endosome integrity. Molecular biology of the cell 15: 5283–5294

Shin HW, Takatsu H (2019) Substrates of P4-ATPases: beyond aminophospholipids (phosphatidylserine and phosphatidylethanolamine). FASEB J 33: 3087–3096

Shin HW, Takatsu H (2022) Regulatory Roles of N- and C-Terminal Cytoplasmic Regions of P4-ATPases. Chem Pharm Bull (Tokyo) 70: 524–532

Stauffer TP, Ahn S, Meyer T (1998) Receptor-induced transient reduction in plasma membrane PtdIns(4,5)P2 concentration monitored in living cells. Curr Biol 8: 343–346.

Takada N, Naito T, Inoue T, Nakayama K, Takatsu H, Shin HW (2018) Phospholipid-flipping activity of P4-ATPase drives membrane curvature. EMBO J 37: e97705

Takada N, Takatsu H, Miyano R, Nakayama K, Shin HW (2015) ATP11C mutation is responsible for the defect in phosphatidylserine uptake in UPS-1 cells. Journal of lipid research 56: 2151–2157

Takatsu H, Baba K, Shima T, Umino H, Kato U, Umeda M, Nakayama K, Shin H-W (2011) ATP9B, a P4-ATPase (a Putative Aminophospholipid Translocase), Localizes to the trans-Golgi Network in a CDC50 Protein-independent Manner. J Biol Chem 286: 38159–38167

Takatsu H, Takayama M, Naito T, Takada N, Tsumagari K, Ishihama Y, Nakayama K, Shin HW (2017) Phospholipid flippase ATP11C is endocytosed and downregulated following Ca2+-mediated protein kinase C activation. Nature communications 8: 1423

Takatsu H, Tanaka G, Segawa K, Suzuki J, Nagata S, Nakayama K, Shin HW (2014) Phospholipid Flippase Activities and Substrate Specificities of Human Type IV P-type ATPases Localized to the Plasma Membrane. J Biol Chem 289: 33543–33556

Tanaka Y, Ono N, Shima T, Tanaka G, Katoh Y, Nakayama K, Takatsu H, Shin HW (2016) The phospholipid flippase ATP9A is required for recycling pathway from endosomes to the plasma membrane. Molecular biology of the cell 27: 3883–3893

Thibaud D, Felix K, Michelle Juknaviciute L, Charlott S, Rasmus Kock F, Syma K, Guillaume L, Joseph AL, Kresten L-L, Poul N (2023) Activation and substrate specificity of the human P4-ATPase ATP8B1. bioRxiv: 2023.2004.2012.536557

Tone T, Nakayama K, Takatsu H, Shin HW (2020) ATPase reaction cycle of P4-ATPases affects their transport from the endoplasmic reticulum. FEBS Lett 594: 412–423

van der Velden LM, Stapelbroek JM, Krieger E, van den Berghe PVE, Berger R, Verhulst PM, Holthuis JCM, Houwen RHJ, Klomp LWJ, van de Graaf SFJ (2010a) Folding defects in P-type ATP 8B1 associated with hereditary cholestasis are ameliorated by 4-phenylbutyrate. Hepatology 51: 286–296

van der Velden LM, Wichers CGK, van Breevoort AED, Coleman JA, Molday RS, Berger R, Klomp LWJ, van de Graaf SFJ (2010b) Heteromeric Interactions Required for Abundance and Subcellular Localization of Human CDC50 Proteins and Class 1 P4-ATPases. Journal of Biological Chemistry 285: 40088–40096

Vance JE (2015) Phospholipid synthesis and transport in Mammalian cells. Traffic 16: 1–18

Wang J, Molday LL, Hii T, Coleman JA, Wen T, Andersen JP, Molday RS (2018a) Proteomic Analysis and Functional Characterization of P4-ATPase Phospholipid Flippases from Murine Tissues. Scientific reports 8: 10795

Wang L, Iwasaki Y, Andra KK, Pandey K, Menon AK, Butikofer P (2018b) Scrambling of natural and fluorescently tagged phosphatidylinositol by reconstituted G protein-coupled receptor and TMEM16 scramblases. J Biol Chem 293: 18318–18327

Wang Y, Menon AK, Maki Y, Liu YS, Iwasaki Y, Fujita M, Guerrero PA, Silva DV, Seeberger PH, Murakami Y et al (2022) Genome-wide CRISPR screen reveals CLPTM1L as a lipid scramblase required for efficient glycosylphosphatidylinositol biosynthesis. Proc Natl Acad Sci U S A 119: e2115083119

Zachowski A (1993) Phospholipids in animal eukaryotic membranes: transverse asymmetry and movement. The Biochemical journal 294 (Pt 1): 1–14

Zhang Y, Werling U, Edelmann W (2012) SLiCE: a novel bacterial cell extract-based DNA cloning method. Nucleic acids research 40: e55

